# Overview of biomedical and public health reviews in Ethiopia from 1970 to 2018: trends, methodological qualities, gaps and future directions

**DOI:** 10.1101/405555

**Authors:** Tesfa Dejenie Habtewold, Sisay Mulugeta Alemu, Shimels Hussien Mohammed, Aklilu Endalamaw, Mohammed Akibu Mohammed, Andreas A. Tefera, Abera Kenay Tura, Nigus Gebremedhin Asefa, Balewgizie Sileshi Tegegne

## Abstract

**Introduction:** Globally, there has been a dramatic increment of narrative reviews, systematic reviews and overview publication rates. In Ethiopia, only small number of reviews are published and no overviews conducted in biomedical and public health disciplines. Therefore, we aimed to (1) assess the trend of narrative and systematic reviews in Ethiopia, (2) examine their methodological quality and (3) suggest future directions for improvement.

**Methods:** PubMed, EMBASE, Web of Science, SCOPUS, CINHAL, WHO Global Index Medicus, Cochrane Library and PsycINFO electronic databases were searched and supplemented by hand searching as well. All narrative reviews and systematic reviews with or without a meta-analysis from 1970 to April 2018 were included. The International Narrative Systematic assessment (INSA) for narrative reviews and A MeaSurement Tool to Assess Systematic Reviews (AMSTAR-2) for systematic reviews with or without a meta-analysis were used for quality appraisal. Fisher’s exact test at the p-value threshold of 0.05 was used to compare the differences in methodological quality.

**Results:** Of the 2,201 initially identified articles, 106 articles published from 1970 to 2018 were eligible for full-text review. Among included reviews, 50.9% were narrative reviews, 16% were systematic reviews and 33.1% were systematic reviews with meta-analyses. Twenty-nine percent were published in Ethiopia and 43.4% were published after 2015. 85.1% of narrative reviews poorly described the characteristics of included studies and 63.8% did not report a conflict of interest. In systematic reviews, 89.6%, 91.7%, and 100% did not register/publish the protocol, justifying the selection of the study designs for inclusion and report sources of funding for the primary studies respectively. Overall, 55.3% of narrative reviews and 75% of systematic reviews with or without meta-analysis had poor methodological quality.

**Conclusions:** Although publication rate of narrative and systematic reviews have risen in Ethiopia, half of the narrative reviews and three-quarters of the systematic reviews had poor methodological quality. We recommend authors to strictly follow standardized quality assessment tools during conducting reviews. Moreover, immediate interventions such as providing methodological training and employers, editors and peer-reviewers should carefully evaluate all reviews before submission or publication.

**What is new?:** *Key findings:*

- The publication rate of narrative and systematic reviews have risen in Ethiopia.
- Almost half of narrative reviews and three-fourths of systematic reviews with or without meta-analysis had poor scientific methodological quality.

*What this adds to what is known:*

- To our knowledge, this is the first overview of its kind providing insight into the publication trend of narrative and systematic reviews, and their methodological rigor in Ethiopia.

*What is the implication, what should change now:*

- Our review shows that the methodological quality of reviews in biomedical and public health discipline in Ethiopia is substantially low and urges immediate intervention.
- We recommended authors to strictly follow standardized quality assessment tools during designing, conducting and reporting (systematic)reviews.

## Introduction

Health care research aims to advance scientific knowledge, understand the risk factors of ill health, and support improvements in the prevention and treatment of diseases.^1^ Carefully designed and implemented research has an enormous impact in the development of any nation; poor quality research on the other hand, is devastating and could lead to suboptimal health outcomes.^2^ Health research is increasing exponentially; for instance in 2016, 869,666 biomedical and public health research citation were indexed in MEDLINE.^3^ The increased publication of scientific research has led to the development of new therapies, guidelines, invention of new methodology to combine results from primary studies and remarkable improvements in healthcare decision making.^5, 6^

In the hierarchy of evidence, rigorously conducted systematic reviews and meta-analyses are at the highest rank to correctly inform decision makers.^7^ In the last four decades, systematic reviews and meta-analyses have been published in biomedical and public health disciplines.^8^ In 2014, 8,000 systematic reviews (22 per day) were indexed in MEDLINE.^9^ Whenever a systematic review is impossible, narrative (also known as historic review or scoping review) can be used to synthesize available evidence, exploring the development of particular ideas and for advancing conceptual frameworks.^10^ Currently, 80,000 narrative reviews are being published per year.^11^ However, reviews have a number of methodological challenges including study selection, use of relevant databases and quality assessment.^12, 13^ To use narrative reviews and systematic reviews with or without meta-analysis for a decision making, they should be conducted to a high standard of quality and continuous quality appraisal is relevant.^14^ Thus, the Cochrane Collaboration has proposed an overview of reviews, new type of study to compile multiple evidence from (systematic)reviews into a single document that is accessible and useful.^15, 16^ The number of overviews has increased from 1 in 2000 to 14 in 2010.^17^ Several institutions and methodologists have designed strategies and tools to synthesize and evaluate methodological quality, quality of evidence and implications for practice despite none being exclusively and universally accepted.^16, 18^ There is a tremendous disparity in research given that narrative reviews, systematic reviews and their quality appraisal tools are mostly published in developed countries.^19^ The contribution of researchers from low-income setting, including Ethiopia, to this publication industry is minimal and needs several interventions.

According to 2006 World Health Organization (WHO) survey, Ethiopia is one of the African countries that has health research policy (HRP), functional national health research system (NHRS) and Scientific/Ethical Review Committee (S/ERC), health institutions with institutional review committees (IRC), national health research institute (NHRI) and faculty of health sciences at the national universities.^20, 21^ In Ethiopia, biomedical and public health research is currently conducted by Central Statistics Agency of Ethiopia (CSA), Armauer Hansen Research Institute (AHRI), Ethiopian Public Health Institute (EPHI), Ethiopian Food, Medicine and Health Care Administration (FMHCA), federal and regional government organizations, academic institutions, private research bodies, non-governmental and international organizations, individual researchers and professional associations.^22^ The research mainly focuses on alleviating major existing health problems including infectious diseases, malaria, diarrhea, acute respiratory infections, tuberculosis, malnutrition, HIV/AIDS, and other sexually transmitted illnesses.^22^ Professional associations, and private and governmental academic institutions host annual national and international research conferences aimed at the transfer of knowledge and skill. In the last decade, the number of primary researches, narrative reviews and systematic reviews with or without meta-analysis has been increased. For example, 189 and 157 articles have been published in BioMed Central and Public Library of Science (PLoS) international Open Access journals respectively.^4^

Despite the aforementioned efforts, biomedical and public health research in Ethiopia is still at an early stage. Studies are mainly observational, and the number of health researchers, research institutes and the overall volume of research output are small.^23, 24^ The major challenges are limited budget, brain drain, low attitude and motivation of professionals (i.e. conducting research only for monetary gain and/or for academic career promotion), low research awareness among the public, lack of communication of research output, and infrastructural problems, such as lack of computers, storage devices and internet access.^4, 25^ Cohort based studies are small and not well organized. Currently, the Butajira birth cohort^26, 27^, Ethiopian Demographic and Health Survey (EDHS)^28^ and university-based demographic surveillance sites are the main sources of data for research albeit limited availability of data. Individual researchers and organizational data archival system are poor. Open grant application opportunities are small and the majority of the researches are funded by the government but this source does not suffice.^24^ Universities have not been tracking published researches. Furthermore, many studies are still published in non-peer reviewed and low-impact journals, and/ or by predatory publishers. Moreover, the quantity and quality of reviews are not yet known.

To our knowledge, there is no overview of narrative and systematic reviews in Ethiopia. Therefore, we aimed to (1) assess the trend of narrative and systematic reviews in Ethiopia, (2) examine the methodological quality using a standardized quality appraisal tool, and (3) provide future directions for researchers, research institutes, and healthcare policymakers.

## Methods

### Searching strategy

PubMed, EMBASE, Web of Science, SCOPUS, CINHAL, WHO Global Index Medicus, Cochrane Library and PsycINFO electronic databases were searched to retrieve all published reviews, systematic reviews and/or meta-analyses. We searched for articles that included any combination of the following search terms in their singular or plural form in their title, abstract and keywords: “narrative review”, “historic review”, “review”, “systematic review”, “meta-analysis”, “pooled analysis”, “Ethiopia” and “Ethiop*”. “AND” and “OR” Boolean operators were used to combine terms and develop the search syntax. Search syntax was developed for each database as shown in Table 1. Depending on the indexed term in each database, all search terms might not be used in each database syntax. We have also hand searched the table of contents of Ethiopian Journal of Health Development (EJHD) (1984 to 2018) (http://www.ejhd.org/index.php/ejhd/index), Ethiopian Journal of Health Science (EJHS) (1990 to 2018) (https://www.ju.edu.et/ejhs/), Addis Continental Institute of public health library (2006 to 2018) (http://www.addiscontinental.edu.et/), Ethiopian Journal of Reproductive Health (http://ejrh.org/index.php/ejrh) and Google Scholar. Furthermore, we manually searched gray literature and cross-references of eligible studies.

**Table 1:** Databases and search syntax.

| S.no. | Database | Search syntax |
| --- | --- | --- |
| 1. | PubMed | ((("Review" [Publication Type]) OR "Meta-Analysis" [Publication Type]) AND (((Ethiopia [Mesh] OR Ethiop*)))) |
| 2. | EMBASE | ('Ethiopia'/exp OR Ethiopia OR Ethiop*:ab,ti) AND ('systematic review'/exp OR 'meta analysis'/exp) |
| 3. | Web of science | (TOPIC: (Ethiopia) OR TOPIC: (Ethiop*)) AND (TOPIC: (systematic review) OR TOPIC: (meta analysis)) |
| 4. | SCOPUS | ((TITLE-ABS-KEY(Ethiopia) OR TITLE-ABS-KEY(Ethiop*)) AND (TITLE-ABS-KEY (systematic review) OR TITLE-ABS-KEY(meta analysis))) |
| 5. | CINHAL | ((TX "systematic reviews" OR meta-analysis) AND (TX Ethiopia OR TX Ethiop*)) |
| 6. | WHO Global Index Medicus | (tw:((((Meta-analysis) OR (Systematic review)) AND ((Ethiopia) OR (Ethiop*))))) |
| 7. | Cochrane Database of Systematic Reviews | ((("Ethiopia" or Ethiop*) and ("systematic review" or "meta analysis")):ti,ab,kw |
| 8. | PsycINFO | TI (("systematic review" OR meta-analysis) AND TI (Ethiopia OR ethiop*)) |

### Inclusion and exclusion criteria

All biomedical and public health narrative reviews and systematic reviews with or without meta-analysis in Ethiopia from 1970 to April 2018 were included. Publication types, whether narrative reviews or systematic reviews with or without meta-analysis, were identified based on authors report.^29^ Exclusion criteria were any one of the following: (1) systematic review protocols, (2) both quantitative and qualitative primary studies, (3) regional and international reviews and/or meta-analysis, (4) case reports, case series, commentaries, anonymous reports, duplicate studies and editorials, (5) published in non-English language, (6) articles without full text, (7) review and meta-analysis in non-human subjects and (8) literature reviews following case report. The full texts of all eligible articles were obtained for data extraction.

### Quality appraisal and data extraction

The methodological quality was assessed by two trained reviewers (SM and BS) using the International Narrative Systematic Assessment (INSA)^30^ for narrative reviews and A MeaSurement Tool to Assess Systematic Reviews (AMSTAR-2)^31^ for systematic reviews with or without meta-analyses. INSA has seven items which can be rated as ‘yes’ or ‘no’ (Supplementary file 1). One point was given for each of the following criteria: clarity of background, objective, conclusion and description of selection of studies, study characteristics, result and conflict of interest. A review with a total INSA score of ≥5 points was considered ‘good’ quality review and ‘poor’ otherwise.^30^ INSA is a valid quality assessment tool for narrative reviews.^30^ AMSTAR-2 has 16 items that include various aspects of systematic review and/or meta-analysis (Supplementary file 1). The overall quality of a systematic review and/or meta-analysis was rated as ‘high’, ‘moderate’, ‘low’ and ‘critically low’.^31^ The rating criteria and interpretation has been published elsewhere.^31^ In addition, the following data were extracted: name of the first author, publication year, number of authors, major topic area, publication type (i.e, narrative review, systematic review, systematic review with meta-analysis), publisher, country of publication, volume of journal, affiliation of authors, number of primary studies included, number of databases searched, study design of included studies, quality assessment tool used to assess primary studies, funding (i.e, funded, not funded/non-declared), years of coverage of primary studies search, and protocol registration and/or publication. Affiliation represents the institutional address of all authors when the research was done and categorized as Ethiopian institutions, foreign institutions or both. Whenever difficult to identify publishers and place of publication, the journal web pages were referred. The topic area was defined based on the main outcome variable.

### Statistical analysis

The data were first analyzed descriptively using frequencies and percentages. Fisher’s exact test was used to compare the methodological quality difference in terms of funding, authors affiliation, number of databases searched to access primary studies, number of primary studies included, years of coverage to search primary studies, publisher, and place of publication. All data entry and analyses were carried out by Statistical Package for the Social Sciences (SPSS) version 23.^32^

## Results

### Search results

Totally, 2,162 articles were obtained from searching PubMed (n =806), EMBASE (n =273), Web of Science (n =215), SCOPUS (n =204), CINHAL (n =36), WHO Global Index Medicus (n =624) and Cochrane library (n =4). We could not find any article from PsycINFO. Additionally, 39 articles were found through a hand searching. After removing duplicate articles (n =940), reviews and meta-analyses on non-human subjects (n =73), and non-English articles (n =31), 1,156 articles were ready for title and abstract screening. Of these, 1,050 articles were excluded for various reasons: 718 were regional and international (systematic)reviews and/or meta-analyses, 311 were non-related titles/case reports, 10 were primary studies and 11 were protocols. Therefore, 106 articles were selected for full-text review. Eleven reviews were excluded from quality evaluation due to the absence of full-text; however, we considered them in the descriptive background characteristics of reviews. Finally, the quality of 95 articles (47 narrative reviews and 48 systematic reviews with or without meta-analyses) were appraised. The PRISMA flow diagram of screening and selection process is shown in Fig. 1.

**Figure 1:**
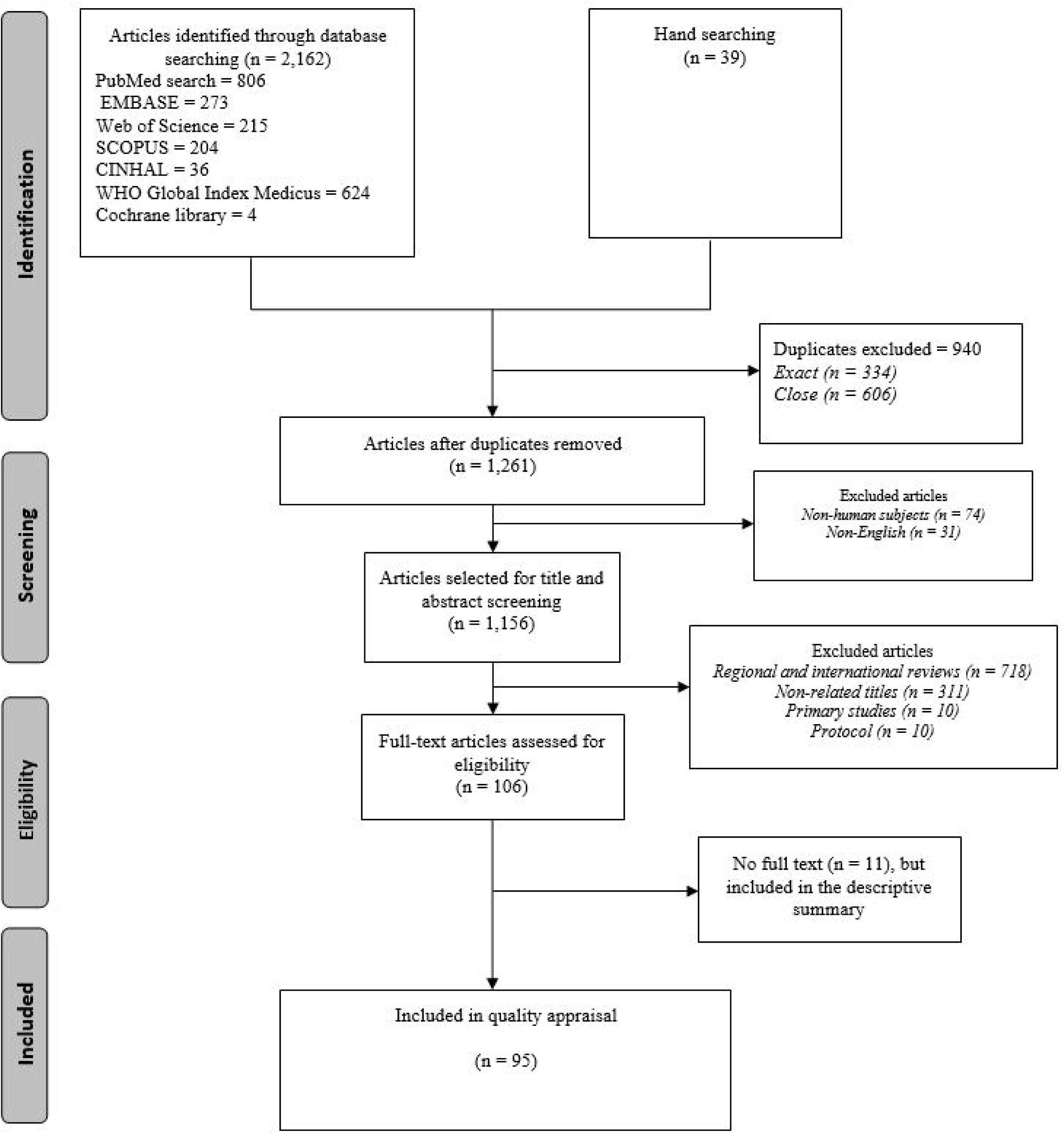
Schematic presentation of literature screening and selection process.

### Background characteristics

Of 106 selected articles published from 1970 to 2018, 54(50.9%) were narrative reviews, 17(16%) were systematic reviews and 35(33.1%) were systematic reviews with meta-analyses. Likewise, 31 (29.2%) were published in Ethiopian journals (e.g. Ethiopian Health Development Journal and Ethiopian Medical Journal), 38(40%) searched three to four databases (i.e, PubMed is the most searched) and 31(32.6%) included 5 to 19 studies. We found only one Cochrane review based on a randomized control trial and only five systematic reviews with meta-analyses have registered their protocol. Table 2 shows summary of included reviews. We also supplemented the detailed characteristics of included reviews (Supplementary file 2).

**Table 2:** Background characteristics of included (systematic)reviews and meta-analyses.

| Characteristics |  | Frequency (n) | Percentage (%) |
| --- | --- | --- | --- |
| Publication year | Before 2011 | 22 | 20.8 |
|  | 2011 to 2015 | 38 | 35.8 |
|  | After 2015 | 46 | 43.4 |
| Number of authors | One | 27 | 25.5 |
|  | 2 to 3 | 40 | 37.7 |
|  | ≥4 | 39 | 36.8 |
| Topic area | HIV/AIDS | 12 | 11.3 |
|  | Nutrition | 10 | 9.4 |
|  | TB | 6 | 5.7 |
|  | Others* | 78 | 73.6 |
| Publication type | Narrative review | 54 | 50.9 |
|  | Systematic review | 17 | 16.0 |
|  | Systematic review with meta-analysis | 35 | 33.1 |
| Publisher | EPHA | 15 | 14.2 |
|  | EMA | 9 | 8.5 |
|  | BioMed Central | 24 | 22.6 |
|  | Jimma University | 7 | 6.6 |
|  | Hindawi | 6 | 5.7 |
|  | PLOS | 6 | 5.7 |
|  | Others | 39 | 43.3 |
| Place of publication | Ethiopia | 31 | 29.2 |
|  | England | 31 | 29.2 |
|  | United States | 22 | 20.8 |
|  | Others | 22 | 20.8 |
| Journal volume | ≤11 | 26 | 24.5 |
|  | 12 to 18 | 28 | 26.4 |
|  | ≥19 | 52 | 49.1 |
| Authors' affiliation(s) | Ethiopia | 69 | 65.1 |
|  | Foreign country | 20 | 18.9 |
|  | Both | 17 | 16 |
| Source databases to access | PubMed | 59 | 55.7 |
| (systematic)reviews and meta-analysis | Google Scholar | 31 | 29.2 |
|  | EMBASE | 6 | 5.7 |
|  | SCOPUS | 6 | 5.7 |
|  | Web of Science | 4 | 3.8 |
| Number of primary studies included<sup>+</sup> | 5 to 19 | 31 | 32.6 |
|  | 20 to 432 | 30 | 31.6 |
|  | Not reported | 34 | 35.8 |
| Number of databases searched to | 1 to 2 | 18 | 18.9 |
| access primary studies<sup>+</sup> | 3 to 4 | 38 | 40.0 |
|  | 5 to 11 | 15 | 15.8 |
|  | Not reported | 24 | 25.3 |
| Study design of primary studies<sup>+</sup> | Observational | 48 | 50.5 |
|  | Randomized control trial | 1 | 1.1 |
|  | Not reported | 46 | 48.4 |
| Protocol registration<sup>++</sup> | Registered | 5 | 10.4 |
|  | Not registered/reported | 43 | 89.6 |
| Quality assessment tool<sup>+</sup> | Standardized tool<sup>-</sup> | 24 | 25.2 |
|  | Not reported/own tool | 71 | 74.8 |
| Funding<sup>+</sup> | Funded | 20 | 21.1 |
|  | Not funded/Non declared | 75 | 78.9 |
| Years of coverage of primary studies | 2 to 15 | 23 | 24.2 |
| search<sup>+</sup> | 15.1 to 72.33 | 23 | 24.2 |
|  | Not reported | 49 | 51.6 |

*\*At least one publication in various topics*
*<sup>+</sup> n=95*
*<sup>++</sup> n=48*
*□ Critical Appraisal for Research, Downs and Black checklists, Evers Checklist, Health states Quality scale, Jadad scale, STrengthening the Reporting of OBservational studies in Epidemiology (STROBE) statement, Joanna Briggs Institute scale, Meta-Analysis of Statistics Assessment and Review Instrument and Newcastle-Ottawa Scale (NOS)* *EPHA=Ethiopian public health association* *EMA=Ethiopian medical association* *PLOS=Public library of science*

Overall, the publication rates of reviews have risen in Ethiopia. The number of a systematic review with or without meta-analysis being published between 2016 and 2018 was four times higher compared with the publication rate between 2011 and 2015. The publication rate of narrative reviews decreased after 2011 while systematic review publication rate remained stable (Fig. 2).

**Figure 2:**
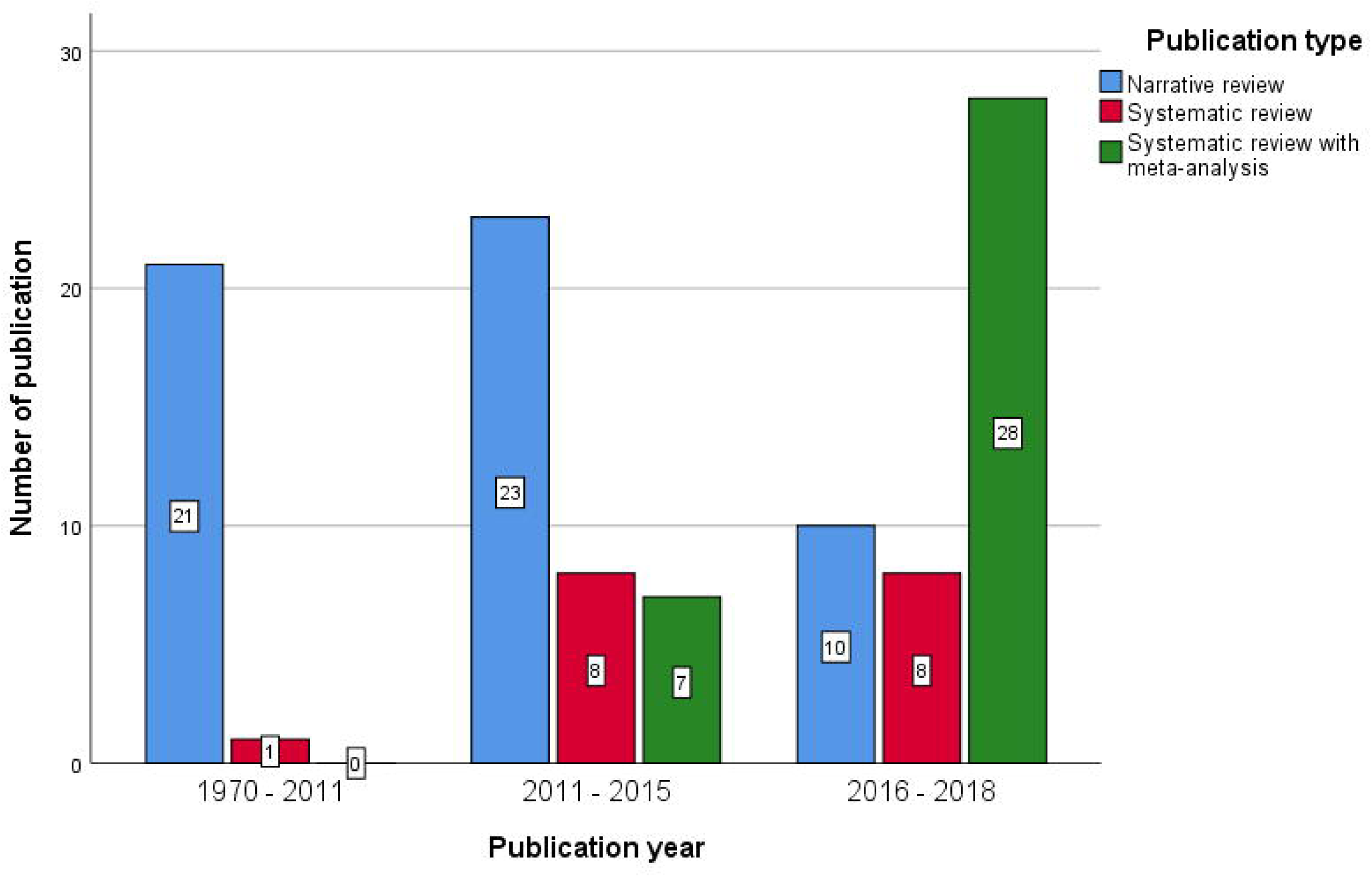
Frequency distribution of (systematic)reviews and meta-analyses publication from 1970 to 2018.

### Methodological quality of narrative reviews

As presented in Table 3, 85.1% of narrative reviews did not clearly describe characteristics of included studies, 63.8% did not report a conflict of interest and 61.7% did not describe the selection of studies. Overall, 26 (55.3%) of narrative reviews had poor quality.

**Table 3:** Description of narrative reviews based on INSA criteria.

| INSA criteria | Yes (n/%) | No (n/%) |
| --- | --- | --- |
| 1. Background of the study clearly explained | 47(100) | 0(0.0) |
| 2. Objective is clear | 37(78.7) | 10(21.3) |
| 3. Description/Motivation of selection of studies | 18(38.3) | 29(61.7) |
| 4. Description of study the characteristics included is clear | 7(14.9) | 40(85.1) |
| 5. Presentation of results (paragraphs, tables, synthesizing of data) | 41(87.2) | 6(12.8) |
| 6. Conclusion is clear | 38(80.9) | 9(19.1) |
| 7. Conflict of interest is stated | 17(36.2) | 30(63.8) |

### Methodological quality of systematic reviews with or without meta-analysis

As shown in table 4, 89.6% of reviews were published without a registered protocol, 91.7% did not justify the selection of the study design for inclusion and none of them reported sources of funding for the primary studies. Overall, 4(8.3%) reviews had high quality, 8(16.7%) had low quality and 36(75%) had critically low quality.

**Table 4:**
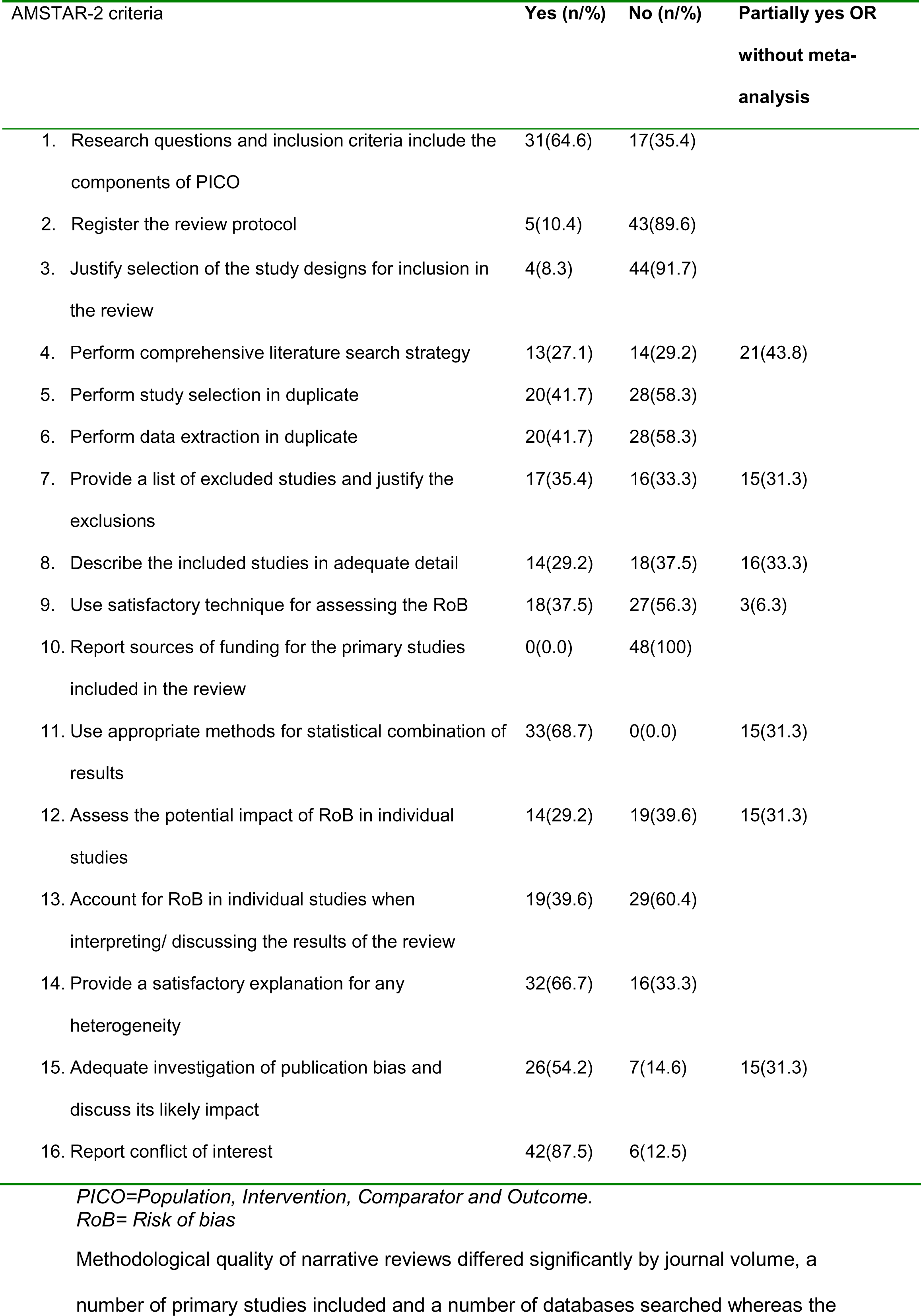
Description of systematic reviews with or without meta-analyses based on AMSTAR-2 criteria

| AMSTAR-2 criteria | Yes (n/%) | No (n/%) | Partially yes OR<br>without meta-analysis |
| --- | --- | --- | --- |
| 1. Research questions and inclusion criteria include the components of PICO | 31(64.6) | 17(35.4) |  |
| 2. Register the review protocol | 5(10.4) | 43(89.6) |  |
| 3. Justify selection of the study designs for inclusion in the review | 4(8.3) | 44(91.7) |  |
| 4. Perform comprehensive literature search strategy | 13(27.1) | 14(29.2) | 21(43.8) |
| 5. Perform study selection in duplicate | 20(41.7) | 28(58.3) |  |
| 6. Perform data extraction in duplicate | 20(41.7) | 28(58.3) |  |
| 7. Provide a list of excluded studies and justify the exclusions | 17(35.4) | 16(33.3) | 15(31.3) |
| 8. Describe the included studies in adequate detail | 14(29.2) | 18(37.5) | 16(33.3) |
| 9. Use satisfactory technique for assessing the RoB | 18(37.5) | 27(56.3) | 3(6.3) |
| 10. Report sources of funding for the primary studies included in the review | 0(0.0) | 48(100) |  |
| 11. Use appropriate methods for statistical combination of results | 33(68.7) | 0(0.0) | 15(31.3) |
| 12. Assess the potential impact of RoB in individual studies | 14(29.2) | 19(39.6) | 15(31.3) |
| 13. Account for RoB in individual studies when interpreting/ discussing the results of the review | 19(39.6) | 29(60.4) |  |
| 14. Provide a satisfactory explanation for any heterogeneity | 32(66.7) | 16(33.3) |  |
| 15. Adequate investigation of publication bias and discuss its likely impact | 26(54.2) | 7(14.6) | 15(31.3) |
| 16. Report conflict of interest | 42(87.5) | 6(12.5) |  |

*PICO=Population, Intervention, Comparator and Outcome.*
*RoB= Risk of bias*

Methodological quality of narrative reviews differed significantly by journal volume, a number of primary studies included and a number of databases searched whereas the quality of systematic reviews with or without meta-analyses differed by use of standardized methodological quality assessment tools (Table 5).

**Table 5:** Differences in methodological quality of narrative review and systematic review with or without meta-analyses.

| Characteristics | Category | Quality of narrative review |  |  | Quality of systematic review |  |  |  |
| --- | --- | --- | --- | --- | --- | --- | --- | --- |
|  |  | Poor | Good | p-value | High | Low | Critically low | p-value |
| Publication year | Before 2011 | 11 | 4 | 0.14 | - | - | - | 0.25 |
|  | 2011-2015 | 9 | 13 |  | 0 | 1 | 13 |  |
|  | After 2015 | 6 | 4 |  | 4 | 7 | 23 |  |
| Number of authors | One | 7 | 3 | 0.25 | 0 | 1 | 11 | 0.09 |
|  | 2 to 3 | 12 | 7 |  | 0 | 2 | 14 |  |
|  | ≥4 | 7 | 11 |  | 4 | 5 | 11 |  |
| Topic area | HIV/AIDS | 3 | 2 | 0.19 | 2 | 1 | 1 | 0.15 |
|  | Nutrition | 3 | 0 |  | 0 | 1 | 5 |  |
|  | TB | 0 | 2 |  | 0 | 0 | 3 |  |
|  | Others* | 20 | 17 |  | 2 | 6 | 27 |  |
| Publisher | EPHA | 9 | 3 | 0.12 | 0 | 0 | 3 | 0.08 |
|  | EMA | 2 | 0 |  | - | - | - |  |
|  | BioMed Central | 0 | 2 |  | 0 | 3 | 18 |  |
|  | Jimma University | 2 | 4 |  | 0 | 1 | 0 |  |
|  | Hindawi | 0 | 1 |  | 2 | 0 | 3 |  |
|  | Others | 13 | 11 |  | 2 | 4 | 12 |  |
| Place of publication | Ethiopia | 13 | 7 | 0.67 | 0 | 1 | 3 | 0.27 |
|  | England | 4 | 3 |  | 0 | 4 | 18 |  |
|  | United States | 4 | 5 |  | 3 | 2 | 7 |  |
|  | Others | 5 | 6 |  | 1 | 1 | 8 |  |
| Journal volume | ≤11 | 4 | 8 | 0.02 | 0 | 1 | 12 | 0.45 |
|  | 12-18 | 1 | 4 |  | 3 | 5 | 13 |  |
|  | ≥19 | 21 | 9 |  | 1 | 2 | 11 |  |
| Authors' affiliation(s) | Both | 4 | 6 | 0.45 | 2 | 2 | 3 | 0.09 |
|  | Foreign country | 17 | 10 |  | 0 | 2 | 5 |  |
|  | Ethiopia | 5 | 5 |  | 2 | 4 | 28 |  |
| Source databases | EMBASE | 1 | 0 | 0.42 | 0 | 1 | 4 | 0.81 |
| access | Google Scholar | 11 | 5 |  | 3 | 3 | 9 |  |
| (systematic)review and | PubMed | 13 | 14 |  | 1 | 4 | 18 |  |
| meta-analysis | SCOPUS | 1 | 2 |  | 0 | 0 | 3 |  |
|  | Web of Science | - | - |  | 0 | 0 | 2 |  |
| Number of primary | 5 to 19 | 0 | 8 | 0.001 | 3 | 3 | 17 | 0.66 |
| studies included<sup>+</sup> | 20 to 432 | 4 | 2 |  | 1 | 5 | 18 |  |
|  | Not reported | 22 | 11 |  | 0 | 0 | 1 |  |
| Number of databases | 1 to 2 | 0 | 5 | <0.001 | 1 | 1 | 11 | 0.11 |
| searched <sup>+</sup> | 3 to 4 | 4 | 10 |  | 0 | 6 | 18 |  |
|  | 5 to 11 | 2 | 3 |  | 3 | 1 | 6 |  |
|  | Not reported | 20 | 3 |  | 0 | 0 | 1 |  |
| Funding<sup>+</sup> | Funded | 7 | 9 | 0.36 | 0 | 0 | 4 | 0.99 |
|  | Not funded/Non-declared | 19 | 12 |  | 4 | 8 | 32 |  |
| Years of coverage of | 2 to 15 | 1 | 2 | 0.08 | 4 | 5 | 11 | 0.08 |
| primary studies search<sup>+</sup> | 15.1 to 72.33 | 3 | 7 |  | 0 | 1 | 12 |  |
|  | Not reported | 22 | 12 |  | 0 | 2 | 13 |  |
| Tools used to assess | Standardized tool<sup>-</sup> |  |  |  | 0 | 0 | 26 | <0.001 |
| methodological quality<sup>+</sup> | Not reported |  |  |  | 4 | 8 | 10 |  |

*\*At least one publication in various topics*
*<sup>+</sup> total number of article is 95*
*□ Critical Appraisal for Research, Downs and Black checklists, Evers Checklist, Health states Quality scale, Jadad scale, STROBE statement, Joanna Briggs Institute scale, Meta-Analysis of Statistics Assessment and Review Instrument and NOS* *EPHA=Ethiopian Public Health Association* *EMA: Ethiopian Medical Association*

## Discussion

This overview, which is the first of its type in Ethiopia, aimed to synthesize evidence on narrative reviews and systematic review with or without meta-analyses. There has been an increasing publication rate of reviews where most have been published in the last decade and were based on basic research. However, half of the narrative reviews and three-quarters of the systematic reviews had a poor methodological quality, which diminishes the trustworthiness, accuracy, and comprehensiveness of newly generated evidence. We also uncovered that the methodological quality of narrative reviews differed by journal volume, a number of primary studies included, and number of databases searched whereas the quality of systematic reviews differed by use of standardized methodological quality assessment tool.

Since 2011, the publication rate of systematic review with or without meta-analysis in Ethiopia has risen fourfold. This might be due to an expansion of higher education institutions, use of publications for academic career promotion and other incentives, increment of publication rate of primary studies, increased budget allocation for research, opportunity for open access waiver fund by publishers and increased collaboration of researchers compared to previous years. The publication rate of narrative reviews has decreased by half since 2011, which might be due to methodological advancement and increased authors’ knowledge about statistical techniques to combine primary studies results.^10^

This overview revealed that almost all narrative and systematic reviews are based on observational studies, and 55.3% of narrative reviews and 75% of systematic reviews with or without meta-analysis had poor scientific methodological quality. Our finding is in agreement with previous overview reports in biomedical and public health outcomes which revealed the methodological quality of reviews is not as high as the publication rate.^33^-^36^ This could be attributed to conducting and reporting systematic reviews and meta-analyses without registering or publishing a study protocol; consequently, authors may be biased. This hypothesis is supported by our findings which shows only 4 out of 48 systematic reviews with or without meta-analyses have registered their protocol in the International Prospective Register of Systematic Reviews (PROSPERO). This problem can be supplemented by the publication of these reviews in low impact journals, which are less likely to request the registration of a protocol before considering the reviews for publication. In 2016, two-thirds of published protocols have been registered in PROSPERO^37^; systematic reviews with registered protocols have high quality compared with reviews without registered protocols.^38^ Another possible explanation is that authors may not be aware of the quality criteria given that most tools are recently invented.^18, 30, 39^-^41^ Inadequate quality assessment of included primary studies using a standardized tool may also explain our findings.^19^ For example, only one-fourth of the reviews assessed the methodological quality of primary studies using a standardized tool, such as Joanna Briggs Institute (JBI) critical appraisal tools, Downs and Black checklists, Evers Checklist, Jadad scale, STrengthening the Reporting of OBservational studies in Epidemiology (STROBE) statement, Meta-Analysis of Statistics Assessment and Review Instrument (MAStARI) and Newcastle-Ottawa Scale (NOS). In addition, peer reviewers, editors and authors may not be fully aware of meta-analysis methodological approaches and can be unqualified to address statistical issues.^10^ Involving only one reviewer to screen studies, conducting review without adequate expertise or consulting experts, including small number of studies and faster completion with poorer reporting quality could be possible reasons.^42^ Furthermore, lack of well-established and harmonized criteria for academic carrier promotion and incentivization, which may lead researchers to focus only on the number of publication instead of ensuring the quality of reviews.

In this overview, we summarized evidence from reviews and meta-analyses and evaluated the methodological qualities using a standardized tool. Since this is the first overview of its kind, it also provides an insight into the trend of narrative and systematic reviews and the level of quality of evidence generated in Ethiopia. The difference in scientific methodological quality has been compared in terms of several factors. Our overview is the most comprehensive overview by including all reviews and meta-analyses published so far in biomedical and public health discipline in Ethiopia and provide a nationally representative evidence. We assessed and reported the quality of narrative reviews for the first time using a standardized tool. It is also relevant to acknowledge the limitation of this overview. INSA is the only quality assessment tool for narrative reviews which is broad that can lead to subjective bias. Despite the popularity of AMSTAR, it is also relevant to admit that the revised AMSTAR-2 tool was not validated.^31^ Given the limited number of reviews, only a few public health and biomedical topics were addressed; as a result, topic-based in-depth analysis was not carried out and it may be difficult to translate the result to a particular healthcare intervention or outcome. Furthermore, we could not ascertain a strong association due to small sample size and results should be interpreted with caution.

### Conclusions, implications and future directions

Publication rate of narrative reviews and systematic reviews with or without meta-analysis have risen, but on the other hand, half of the reviews and three-quarters of the systematic reviews had poor methodological quality. Given that developing countries share common problems, other nations should also assess the publication rate and evaluate the scientific methodological quality of narrative and systematic reviews, identify country-specific gaps and provide problem-centered interventions. Most narrative and systematic reviews are epidemiologic reviews focusing on prevalence and associated factors; hence, reviews based on diagnostic, prognostic and therapeutic research are needed. Even though many quality appraisal tools are available for systematic reviews with or without meta-analysis, more objective and detailed assessment tool is required for narrative reviews. Further investigation is also required to identify the facilitators and barriers of uptake and quality of narrative and systematic reviews. Despite the low methodological rigor, almost all systematic reviews and meta-analysis followed the PRISMA statement. We recommend authors to strictly follow standardized quality assessment tools during conducting and reporting (systematic)reviews with or without a meta-analysis.^18, 43^

In general, establishment of sustainable and large financial sources for research with a special emphasis on clinical and applied research, strengthening the new established research councils, training authors, professional methodologists, editors and reviewers, supporting professional associations and using experience of Ethiopian diaspora professionals are important to improve the number and quality of biomedical and public health reviews and overviews in Ethiopia.^10, 19, 25^ Although there was no quality difference, in our overview, only one out of five narrative and systematic reviews are funded. Among others, description of characteristics of included studies, reporting of conflict of interest, description of selection of studies, protocol registration, justifying selection of the study designs for inclusion and reporting sources of funding for the primary studies are least implemented components of a review. Furthermore, establishment of a new national library repositories and strengthening the existing ones, continuing research career promotion and increasing incentives provided to researchers based on impartial, harmonized and evidence-based criteria, increasing the number of local journals and ensuring their peer-review quality and increasing the number of research sites, for example, increasing Demographic Surveillance Systems (DSS) would be very important strategies to ensure output and quality of reviews.^23^ Finally, it is helpful to promote development and uptake of narrative reviews and systematic reviews with or without meta-analyses through informing the importance for healthcare policymakers, increasing access to the international journals, giving priority and support for systematic reviews, increasing competency and willingness of researchers to conduct reviews, creating awareness about importance for end-users, and improving the quality, visibility and accessibility of local primary research.^19, 44^

## Declarations of interest

None

### Author Contributions Statement

Conceived and designed the experiments: T.D.

Performed the experiments: S.M., B.S and T.D.

Analyzed and interpreted the data: T.D.

Contributed reagents, materials, analysis tools or data: S.M., B.S and T.D.

Wrote the paper: T.D., S.M., S.H., M.A., A.E., A.A., A.K., N.G., and B.S.

Final approval of the version to be submitted: T.D., S.M., S.H., M.A., A.E., A.A., A.K., N.G., and B.S.

